# Evaluating the ALERT algorithm for local outbreak onset detection in seasonal infectious disease surveillance data

**DOI:** 10.1101/664433

**Authors:** Alexandria C. Brown, Stephen A. Lauer, Christine C. Robinson, Ann-Christine Nyquist, Suchitra Rao, Nicholas G. Reich

## Abstract

Estimation of epidemic onset timing is an important component of controlling the spread of seasonal infectious diseases within community healthcare sites. The Above Local Elevated Respiratory Illness Threshold (ALERT) algorithm uses a threshold-based approach to suggest incidence levels that historically have indicated the transition from endemic to epidemic activity. In this paper, we present the first detailed overview of the computational approach underlying the algorithm. In the motivating example section, we evaluate the performance of ALERT in determining the onset of increased respiratory virus incidence using laboratory testing data from the Children’s Hospital of Colorado. At a threshold of 10 cases per week, ALERT-selected intervention periods performed better than the observed hospital site periods (2004/2005-2012/2013) and a CUSUM method. Additional simulation studies show how data properties may effect ALERT performance on novel data. We found that the conditions under which ALERT showed ideal performance generally included high seasonality and low off-season incidence.

## 1 INTRODUCTION

In healthcare settings, policies that enforce the use of enhanced personal protective equipment are some of the important interventions that can reduce infectious disease spread [1]. The control of seasonal outbreaks within community healthcare institutions is important for public health, particularly in healthcare settings where the young, elderly, and immunocompromised are at the highest risk. One component of infectious disease control is early detection, which is a goal of infectious disease surveillance systems at the local, state, and national levels [2]. This transition from endemic to epidemic activity is critical, as it corresponds to an increase in demand for healthcare and necessitates the implementation of protective measures. During outbreak periods, the “epidemic onset” occurs when case counts rise above a pre-defined background level [3].

Upper respiratory illnesses, caused by influenza A, influenza B or respiratory syncytial virus (RSV), are common in temperate regions worldwide. Seasonal outbreaks of these viruses are one of the primary reasons that healthcare facilities implement periods of time where enhanced infection precautions are enforced [1]. In one study of the mortality associated with these infections, influenza A (H3N2) caused the highest number of deaths, followed by RSV, influenza B, and influenza A (H1N1) viruses [4]. The actual toll of these illnesses is difficult to calculate as upper respiratory viruses are often accompanied by circulatory or pneumonia complications, especially in the young and elderly. Estimates of average annual influenza-related deaths in the United States range from 10,682 to 28,169, while RSV-related average annual deaths have been estimated at 6,211 to 17,199 [5].

In the United States, the Centers for Disease Control and Prevention (CDC) defines the influenza season as beginning in November and ending in April, with influenza activity seen as early as October and as late as May in some regions. The exact dates related to the onset of the influenza season vary at the state and local level.

Many hospital sites currently use either a threshold-based or date-based trigger to signal the onset of respiratory illness season. Selection of these triggers is often based on anecdotal observations based on historical incidence or monitoring of local or regional influenza activity. Practical challenges to implementing seasonal policies may provide motivation for healthcare sites to decrease the duration of these periods as soon as possible while still controlling infection spread. Increased personal protective equipment is expensive and often unpopular among healthcare workers, patients, and visitors, which can be found documented in the personal protective equipment compliance literature [6]. Efficient selection of the intervention periods would be financially savvy for hospitals and clinics while maximizing patient protection. Increased numbers of patients during respiratory season drive the need for hospitals to add additional staff (traveling nurses) to be able to provide safe care. Administration has the challenge of when to bring in temporary support and how long to retain them each year. Predicting the increase in seasonal respiratory illness season would enable them to limit the time for the contract and potentially decrease the expense.

Many approaches have been used to detect and characterize transitions between endemic and epidemic incidence patterns. For the stochastic prediction of infectious disease spread between individuals, mechanistic models (such as agent-based [7] and compartmental susceptible-infectious-recovered [8], among others) are well developed and have been implemented as stand-alone forecasting models [9, 10]. Autoregressive integrated moving average (ARIMA) or seasonal ARIMA (SARIMA) models are well-known statistical approaches for modeling time-series, such as infectious disease case counts, that correlate with past observations [11, 12]. Both statistical and mechanistic models have been used successfully in infectious disease forecasting [13, 14]. However, these methods on their own are not designed specifically to detect onset periods or guide real-world policy. Other research has used thresholds in order to characterize influenza incidence into low, moderate and high categories using a “Moving Epidemic Method”, which uses maximum accumulated rates percentage (MAP) based on incidence rates per 100,000 inhabitants, or consultations [15, 16]. This approach characterizes the intensity of influenza epidemics and can also trigger enhanced protective interventions, but is intended to be used on a larger scale than that available at even the largest hospital sites. The most common algorithms that are designed to trigger interventions during an outbreak are CUSUM-based methods and their variants [17, 18], the exponential weighted moving average (EWMA) [19, 20], and the space–time permutation scan statistic model [21, 22]. All of these methods involve detections of deviations from expected values, or threshold values, based on historical data. These statistical methods may require advanced statistical training and computational resources to implement at the local level.

The Above Local Elevated Respiratory Illness Threshold (ALERT) algorithm [23] uses a threshold-based trigger system to help healthcare workers determine the epidemic onset prior to the start of the outbreak period. It is available online both as a free R software package and a graphical web applet (http://reichlab.github.io/alert.html). More detailed instruction on using the package and application are available in the ALERT package documentation (Appendix 1) (https://github.com/reichlab/ALERT/blob/master/vignettes/ALERTDocumentation.pdf).

Seasonal infectious disease surveillance data often shows regular patterns of onset, peak, and nadir [24]. The goal of ALERT is to assign a static value to the incidence threshold level used to define epidemic onset. Using historical information from a local surveillance system (e.g. a hospital or city), ALERT assists in the choosing of an appropriate time to begin a particular intervention that would cover the period of highest seasonal respiratory virus activity. Prior work has shown how ALERT can assist in determining the timing of hospital-based interventions for influenza [23]. Likewise, ALERT has been used previously to detect the onset of upper respiratory illness season in the Respiratory Protection Effectiveness Clinical Trial (ResPECT, https://clinicaltrials.gov/ct2/show/NCT01249625), a comparison of N95 and medical masks to protect healthcare workers from seasonal viruses. In this work, we provide a technical overview of the ALERT algorithm, a motivating example, and a simulation study in which we characterize ALERT’s performance over datasets with varying parameters derived from the influenza and RSV datasets.

## 2 METHODS

### 2.1 Model Framework

The ALERT algorithm is a tool for triggering epidemic infection control measures that is designed to capture the most epidemic activity while minimizing the duration of the identified period. The algorithm uses local, site-specific historical data to establish a set of threshold case count values to represent the onset of the epidemic season. The “ALERT period” is a window of time between this onset and when the seasonal peak has likely passed. An ALERT period begins when the reported number of laboratory-confirmed cases in single time unit exceeds a given threshold. The ALERT period ends when the reported number of cases falls below that same threshold after a pre-specified minimum amount of time.

Surveillance data is often provided to researchers as case counts ordered across time, as shown in the left panel of figure 1. Let *y*_*s,t*_ denote the number of cases of a single disease or multiple, pooled pathogens observed at a location in season *s*, *s* = 1, …, *S*, at time unit *t*, *t* = 1, …, *T*. For example, if data is aggregated into weekly time units, *T* = 52.

**FIGURE 1:**
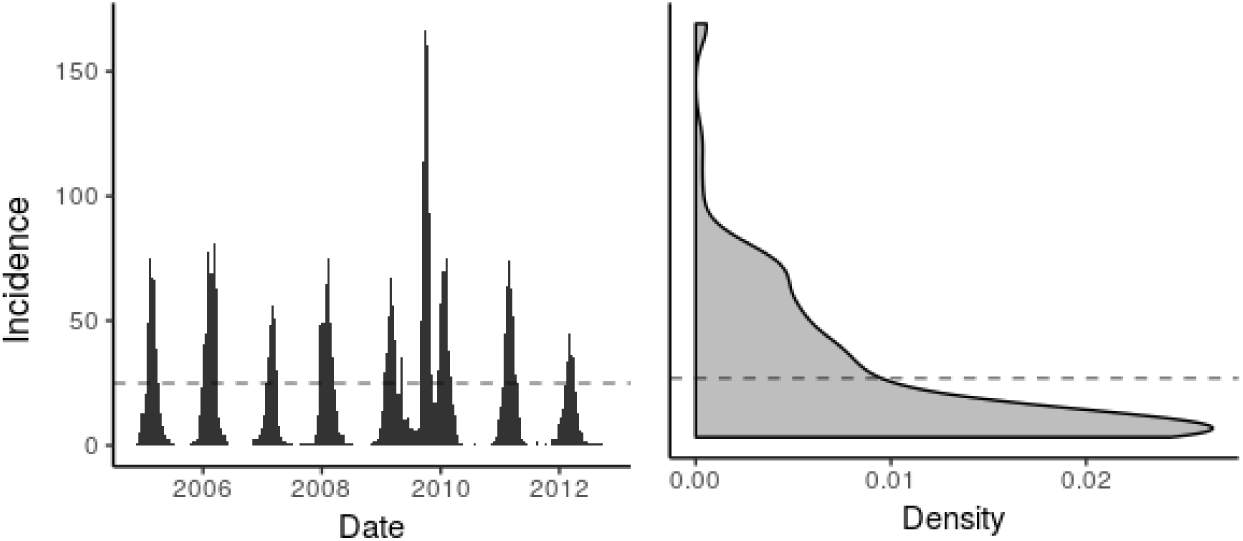
Historical case counts of influenza A, influenza B, and RSV detections combined for the Children’s Hospital of Colorado system are shown at left. A density plot of the case counts in the left panel. In both panels, lab-confirmed respiratory illness incidence is shown on the vertical axis. ALERT calculates percentiles of interest from non-zero cases using a quantile function *Q*_*p*_ (*y*), where the output is the *p*th percentile of *y*. The value of *Q*_*p*_ (*y*), represented in this figure as a hypothetical value of 25 by the dashed line, is selected as a potential epidemic onset threshold, *τ*_*p*_.

Percentiles of interest are calculated from all of the non-zero cases using a quantile function *Q*_*p*_ (*y*), where the output is the *p*th percentile of *y*, as shown in the right panel of figure 1. A value of *Q*_*p*_ (*y*) is selected as a potential epidemic onset threshold, *τ*_*p*_, where *τ*_*p*_ = *Q*_*p*_ (*y*). The set of ***τ*** can be specified as, for instance, all integer thresholds between the 10th and 60th percentile. When incidence exceeds the trigger value *τ*_*p*_, which is selected by the user, this represents the beginning of the ALERT period. The ALERT period extends until both the incidence falls below the trigger and the minimum ALERT duration—also set by the user—has elapsed.

After ***τ*** is determined, the observed *y*_*s,t*_ are used to calculate additional metrics for each *τ*_*p*_. For each *τ*_*p*_, the ALERT algorithm summarizes data from previous years as if that threshold had been applied. If we have historical data on *S* seasons, let *D*_*s*_ be the duration of the ALERT period for season *s* and threshold *τ*_*p*_. *D*_*s*_ is determined by the number of time units (often weeks) from the first instance of *y*_*s,t*_ ≥ *τ*_*p*_ to the following *y*_*s,t*_ ≤ *τ*_*p*_, provided that *D*_*s*_ is larger than a pre-determined minimum duration set by the user. This prevents an ALERT period from being prematurely terminated by early-season fluctuations around the trigger threshold. Additionally, to account for reporting delays and possible delays in implementation of any policies, the user may specify a lag period: a number of time periods between the reporting date associated with the trigger and the date the ALERT period becomes effective. For instance, lag periods may be helpful in tuning ALERT to accommodate a municipal or state health department wanting to implement an increased upper respiratory protection program that needs to wait for reporting from area hospital or requires time to distribute or set up the intervention. Let *X*_*s*_ be the percentage of cases captured in the ALERT period for each season.

The following metrics are calculated and reported to the user for each *τ*:

1. Across all seasons, the median percentage of all cases contained within the ALERT period, *medi an*(*X*_*s*_).
2. Across all seasons, the minimum and maximum of *X*_*s*_.
3. Across all seasons, the median ALERT period duration, *median*(*D*_*s*_).
4. The proportion of seasons in which the ALERT period contained *max*_*t*_ (*y*_*s*_*t*), or the “epidemic peak”, *P C*_*s*_.
5. The proportion of seasons in which the ALERT period contained the peak week ± *k* weeks, where *k* is specified by the user.
6. The mean number of “low weeks” included in the ALERT period; weeks with counts less than *τ*_*p*_, *W C*_*s*_.
7. The mean difference between, for each season, the duration of the ALERT period and the duration of the shortest period needed to capture a pre-determined target percent of cases for that season.

## 3 MOTIVATING EXAMPLE

The datasets used in the following example were provided by the Children’s Hospital of Colorado (CHCO), a 444-bed hospital in Aurora, Colorado serving the Denver metropolitan and surrounding areas. CHCO employs a passive surveillance system where patients with respiratory symptoms are tested for common respiratory viruses at clinicians’ discretion. During the study period, initial methods of virus detection were culture and antigen testing, with polymerase chain reaction used exclusively from 2009 on. In some years, the respiratory cases seen at CHCO after the increased respiratory virus protection policies had already been triggered were not tested, resulting in mid-season under-reporting.

In anticipation of the respiratory virus season each year, CHCO implements additional protective measures for patients and providers including enhanced personal protective equipment (PPE) and restrictions in the visitor policy. The periods of increased upper respiratory protection used historically by CHCO varied across the dataset used in this paper. The initial trigger to declare increased interventions in response to seasonal increased upper respiratory virus incidence was 3 or more lab confirmed cases per day. Later CHCO switched to a date-based system (Dec 1-April 30) derived from anecdotal local incidence patterns.

We compared the ALERT algorithm to a CUSUM baseline model for a portion of the combined RSV, influenza A, and influenza B data from the respiratory virus incidence starting in 2004 to 2012. We chose the 2004/2005-2012/2013 subset of the full data (2002/2003-2012/2013) because data on the time periods of increased protection implemented at the hospital site are not available prior to the 2004 season.

The dataset was divided into training seasons (2004/2005-2007/2008) and testing seasons (2008/2009-2011/2012). We applied the ALERT algorithm to the training set to choose a set of threshold values and compared those to hospital-derived PPE periods, results of which are shown in Table 1. The CUSUM approach was calibrated to give a median (*D*_*s*_) of 19.0 weeks, to match the observed intervention periods. In the training portion of the dataset, the hospital-derived increased PPE periods lasted a median (*D*_*s*_) of 19.0 weeks, ranging from 20.0 weeks in the 2005/2006 and 2006/2007 seasons to 18.0 weeks in 2007/2008 and 2008/2009. We chose trigger thresholds of 6, 10, and 21 cases as illustrative examples, as shown in Table 1 and Figure 2. Notably, *τ*_*p*_ = 10 results in a median *D*_*s*_ that is equivalent to the median *D*_*s*_ of the hospital-derived respiratory protection period.

**TABLE 1:**
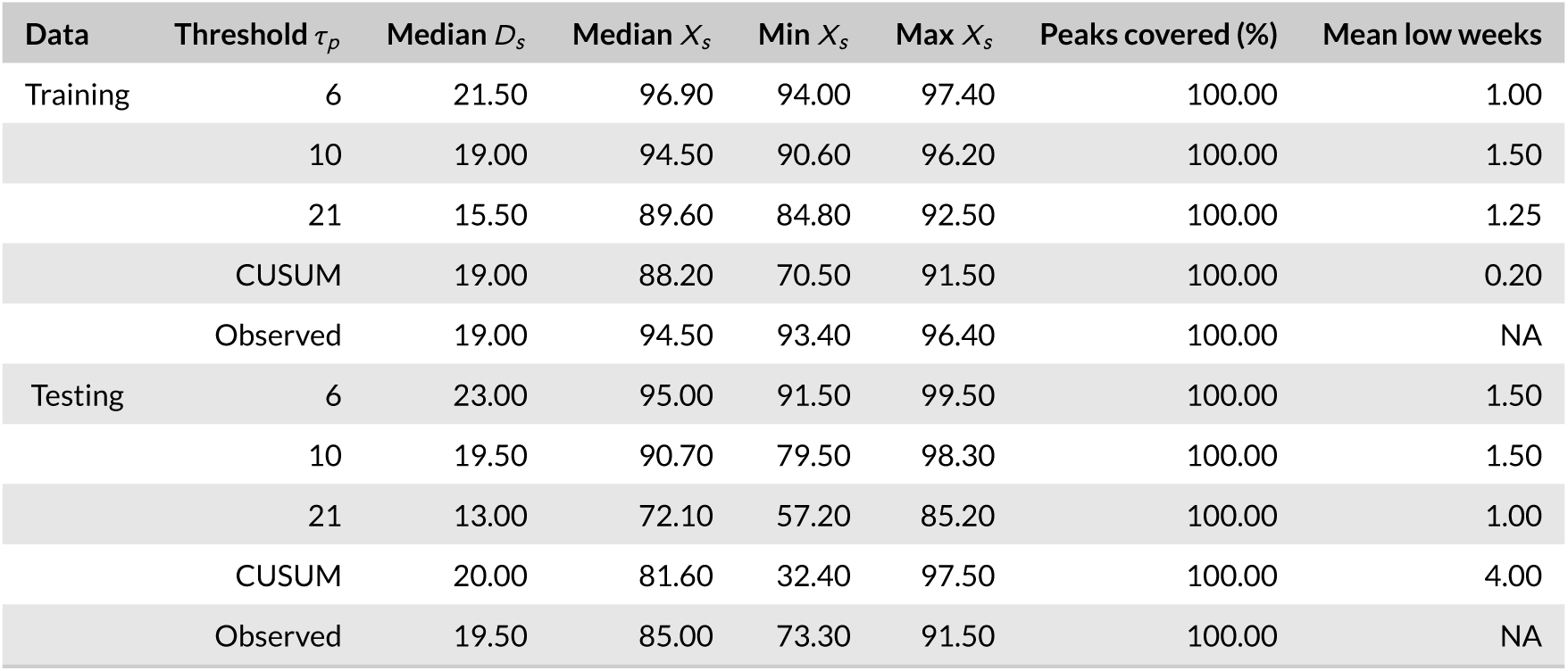
ALERT performance on combined Children’s Hospital of Colorado RSV, influenza A, and influenza B case data for the training (2004/2005-2008/2009) and testing (2009/2010-2012/2013) portions of the dataset. We compared ALERT’s performance at 3 different thresholds to the intervention periods used at the hospital site and a CUSUM approach. For each threshold (*τ*_*p*_), ALERT calculates the median duration (*D*_*s*_), the median, minimum, and maximum percentage of cases covered (*X*_*s*_), the percentage of peaks covered, and mean number of weeks below the threshold (mean low weeks). The observed intervention period triggers varied among seasons in the hospital dataset, as described in the text. As these were determined sometimes based on case numbers and sometimes on date-based cutoffs, a meaningful mean low weeks included in the intervention period for the observed data was not calculable.

**FIGURE 2:**
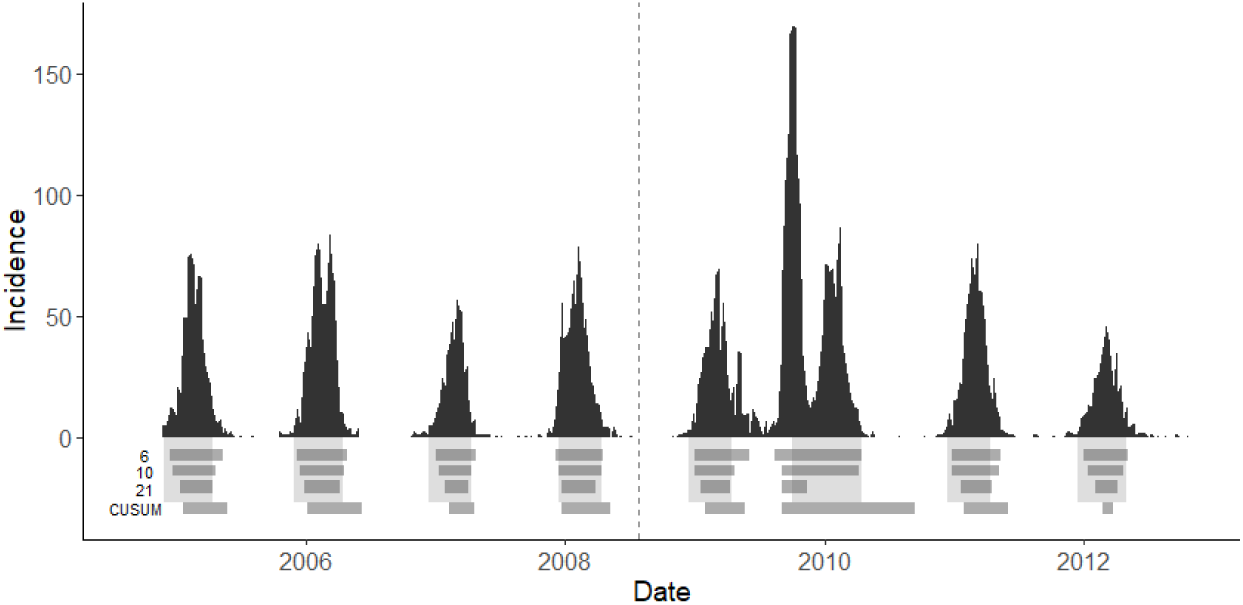
Each panel shows the weekly case counts of combined influenza A, influenza B, and RSV from the Children’s Hospital of Colorado (CHCO) from 2004/2005-2012/2013. Cases to the left of the vertical dashed line were used as the training set for this example, while the testing set appears on the right side of the dashed line. The light gray blocks below the bar graph represent the dates when CHCO implemented increased respiratory protection measures. These periods had a median *D*_*s*_ of 19.5 weeks, with median *X*_*s*_ of 92.5%. The darker horizontal bars show the periods that ALERT would have determined based on the threshold of 6, 10, and 21 cases, or the CUSUM method. Applied to the full dataset, thresholds of 6, 10, and 21 cases yields a median ALERT *D*_*s*_ of 22, 19, and 14 weeks, respectively, with median *X*_*s*_ of 96.9%, 94.5%, and 85.0%. ALERT captures the peak of the 2009 H1N1 outbreak for all thresholds, but ends too early to capture the non-H1N1 seasonal outbreak at *τ*_*s*_ = 21.

The testing portion of the dataset shows that CHCO would have benefited from implementing the ALERT algorithm in their hospital from 2009-2012 at *τ*_*s*_ = 10. Across all years, the sum of all ALERT periods was 145 weeks long, while the as-implemented intervention time totaled 155.4 weeks long. The ALERT period would have covered 5.7% more patients over an equivalent median *D*_*s*_, and saved the hospital a total of 10.4 weeks of intervention time. Furthermore, ALERT would have triggered during the onset of the H1N1 epidemic in 2009, demonstrating that it provides useful information both in seasonal outbreaks settings as well as anomalous pandemic scenarios. Neither *τ*_*s*_ = 6 or *τ*_*s*_ = 21 offered a clear benefit over the periods that were implemented in reality. When *τ*_*s*_ = 6, the median *X*_*s*_ was 95%, but with a concomitant increase in median *D*_*s*_ by 3.5 weeks. Conversely, when *τ*_*s*_ = 21, the median *D*_*s*_ was 6.5 weeks shorter than the observed periods, but yielded a 15.2% decrease in *X*_*s*_. All the ALERT *τ*_*s*_ and that of the observed periods covered 100% of the seasonal peaks. The CUSUM method was calibrated on the training dataset to produce a median intervention period of 19 weeks, similar to the ALERT and observational methods. In both the training and testing datasets, CUSUM did not capture as many cases as the other two methods, however, in the training dataset CUSUM captured the smallest number of low weeks. This did not translate to the testing dataset, however. CUSUM captured the highest number of low weeks in the testing data, and the median *X*_*s*_ was the highest of all the methods. A different CUSUM approach might have performed better than the general approach, but comparison of multiple specialized CUSUM methodologies to find the best fit was outside the scope of this work.

## SIMULATION STUDY

In order to characterize the performance of the ALERT algorithm on time series datasets with varying features, we implemented a simulation study. Using a statistical framework developed by Held, Höhle, and Hofmann, we decomposed the CHCO dataset into a model with parameters that represent known mechanistic attributes, which were varied across a gradient and used to produce many simulated infectious disease datasets [11]. We used the R package [25], surveillance [26] for the estimation and simulation.

We simulated surveillance data using an autoregressive negative binomial model with endemic seasonality. The mean disease incidence *µ*_*t*_ contains autoregressive and endemic components *λ* and *v*_*t*_, respectively, which are modeled as

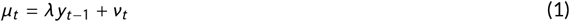

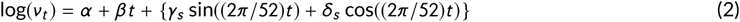

where *λ* > 0 and *v*_*t*_ > 0. In equation 2, *α* is an intercept, *β* is a long-term linear trend parameter, and the bracketed terms represent seasonal variation. *S* is the number of harmonics used (in this case, 1). *γ* and *δ* are parameters that affect noise and season length and timing. *ψ* is an overdispersion parameter which increases the conditional variance of *µ*_*t*_ to *µ*_*t*_ (1 + *µ*_*t*_ *ψ*) for *ψ* > 0.

The range of parameter values that we chose for our simulation study is based on those observed in the CHCO dataset. First, we estimated the mean and standard error for the model parameters for the combined RSV, influenza A and B dataset (Table 2). Second, we set the maximum and minimum simulation values by adding or subtracting twice the standard error from the estimated value for each parameter. Third, we selected 50 evenly-spaced values within this interval. Fourth, for each parameter, we produced 20 simulated time series for each of the 50 values while holding the other parameters constant at their point estimate, resulting in 1000 simulated time series per parameter.

**TABLE 2:** Estimated mean and standard error for the model parameters for the observed weekly case counts from the Children’s Hospital of Colorado data. Parameters are defined in additional detail in equation 2.

| Parameter | Estimate | Standard Error |
| --- | --- | --- |
| $\lambda$ (autoregressive component) | -0.2233 | 0.0577 |
| $\alpha$ (intercept) | 0.2134 | 0.1853 |
| $\beta$ (slope) | -0.0009 | 0.0007 |
| $\gamma$ (noise) | -1.0733 | 0.1654 |
| $\delta$ (season length) | -2.0676 | 0.1518 |
| $\psi$ (overdispersion parameter) | 1.1497 | 0.1255 |

**TABLE 3:** Summary median value of each metric for each parameter. Measures were derived from training and testing datasets by simulation parameter and presented as the median [minimum, maximum] of ALERT duration (*D*_*s*_) and percentage of cases captured (*X*_*s*_).

|  | training |  | testing |  |
| --- | --- | --- | --- | --- |
| | $D_s$ | $X_s$ | $D_s$ | $X_s$ |
| $\alpha$ | 21.0 [11.0, 40.0] | 90.5 [42.4, 99.8] | 19.5 [8.5, 31.0] | 87.7 [46.5, 99.0] |
| $\beta$ | 21.0 [10.0, 41.5] | 90.6 [60.2, 99.7] | 19.5 [8.5, 31.0] | 87.7 [46.5, 99.0] |
| $\delta$ | 21.0 [8.0, 38.0] | 90.8 [18.1, 99.8] | 19.5 [8.5, 31.0] | 88.3 [46.5, 99.0] |
| $\gamma$ | 21.0 [8.0, 38.0] | 90.6 [31.1, 99.7] | 19.5 [8.5, 31.0] | 87.7 [46.5, 99.0] |
| $\lambda$ | 22.0 [8.0, 42.0] | 90.7 [35.4, 99.7] | 19.5 [8.5, 31.0] | 88.3 [46.5, 99.0] |
| $\psi$ | 21.0 [8.0, 43.0] | 91.1 [36.3, 99.5] | 19.5 [8.5, 31.0] | 87.7 [46.5, 99.0] |
| Overall | 21.0 [8.0, 43.0] | 90.7 [18.1, 99.8] | 19.5 [8.5, 31.0] | 87.7 [46.5, 99.0] |

**TABLE 4:** Performance measures ([minimum, maximum]) on training and testing datasets by simulation parameter showing median low weeks captured (*W C*_*s*_) during the ALERT period and median percentage of seasonal peaks captured (*P C*_*s*_).

|  | training |  | testing |  |
| --- | --- | --- | --- | --- |
| | $WC_s$ | $PC_s$ | $WC_s$ | $PC_s$ |
| $\alpha$ | 2.4 [1.0, 5.8] | 80.0 [60.0, 100.0] | 2.5 [1.8, 5.2] | 83.3 [50.0, 100.0] |
| $\beta$ | 2.4 [1.0, 5.4] | 80.0 [60.0, 100.0] | 2.3 [1.8, 5.3] | 83.3 [50.0, 100.0] |
| $\delta$ | 2.4 [1.0, 5.4] | 80.0 [40.0, 100.0] | 2.3 [1.8, 5.3] | 83.3 [50.0, 100.0] |
| $\gamma$ | 2.4 [1.0, 5.4] | 80.0 [40.0, 100.0] | 2.3 [1.8, 5.2] | 83.3 [50.0, 100.0] |
| $\lambda$ | 2.6 [1.0, 5.0] | 80.0 [40.0, 100.0] | 2.3 [1.8, 5.3] | 83.3 [50.0, 100.0] |
| $\psi$ | 2.4 [1.0, 5.2] | 80.0 [60.0, 100.0] | 2.3 [1.8, 5.3] | 83.3 [50.0, 100.0] |
| Overall | 2.4 [1.0, 5.8] | 80.0 [40.0, 100.0] | 2.4 [1.8, 5.3] | 83.3 [50.0, 100.0] |

We set up our simulation study to approximate how the ALERT algorithm would perform in practice; by first tuning the parameters based on some previously observed data, then evaluating the algorithm’s performance on the remainder of the time series. For each time series, the first 260 weeks were used as the training dataset, while weeks 261 through 590 were reserved as the testing dataset. Performance could likely be improved by training on more years of data, however, we chose 5 years to represent a realistic duration of historical data that may be available to many healthcare facilities. ALERT was applied to each training dataset to determine tuning parameters required by the algorithm that would, in practice, be supplied by a knowledgeable clinician or epidemiologist. Ultimately, performance was evaluated by comparing the performance of ALERT in the training phase to that in the testing phase. If there is little change between training and testing phase this implies that the ALERT algorithm can be used to reliably estimate the metrics of interest prospectively.

The best-performing threshold in the training set was determined to be the threshold with the shortest median duration (*D*_*s*_) whose median cases captured (*X*_*s*_) was greater than 85%. The best-performing training threshold was then applied to the testing dataset, which comprised 5 years of simulated data. From these simulations we are able to evaluate the performance of the ALERT algorithm across the values of each parameter. Example simulated datasets for each parameter are represented in Figure 3.

**FIGURE 3:**
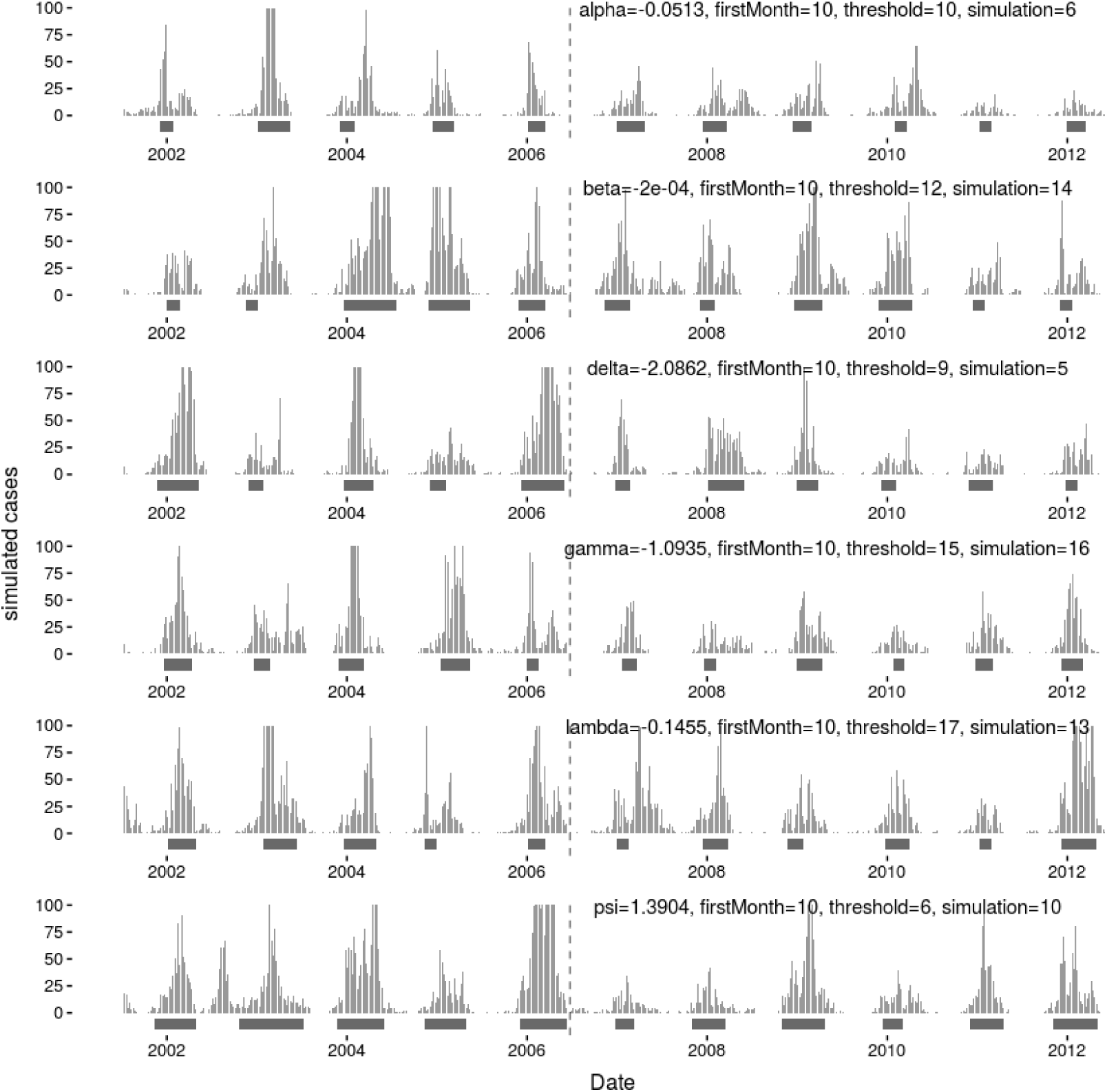
Example simulated time series (vertical bars) for each parameter and the ALERT periods (horizontal bars) corresponding to the threshold that had the median shortest ALERT duration that captured more than 85% of cases during the training set. The vertical dashed line shows the division between the training dataset (left of the line) and testing dataset (right of the line).

ALERT’s performance was remarkably consistent across training and testing datasets for all 4 of the metrics we present here. Across all of the parameters, *D*_*s*_ decreased by 1.5 weeks in the testing datasets, with a concomitant decrease in *X*_*s*_ by 3%. There was no difference in training and testing performance scores in terms of *W C*_*s*_, while *P C*_*s*_ increased for the testing data by 3.3%.

Variations in parameters *α*, *δ*, and *γ* had the greatest impact on ALERT performance metrics. ALERT was less sensitive to changes in *ψ, β*, and *λ*. Figure 4 shows that the percent peaks captured remained more than 76% even in the poorest performers. Increasing values for *α* decreased the percent cases captured by about 10%, while increasing *δ* had the opposite effect. Neither *β* nor *λ* had a marked impact on whether or not seasonal peaks were captured across the range of parameter values that were tested. For the remaining parameters, *γ* and *ψ*, parameter values had a variable impact on percentage of peaks captured, with no obvious linear trend.

**FIGURE 4:**
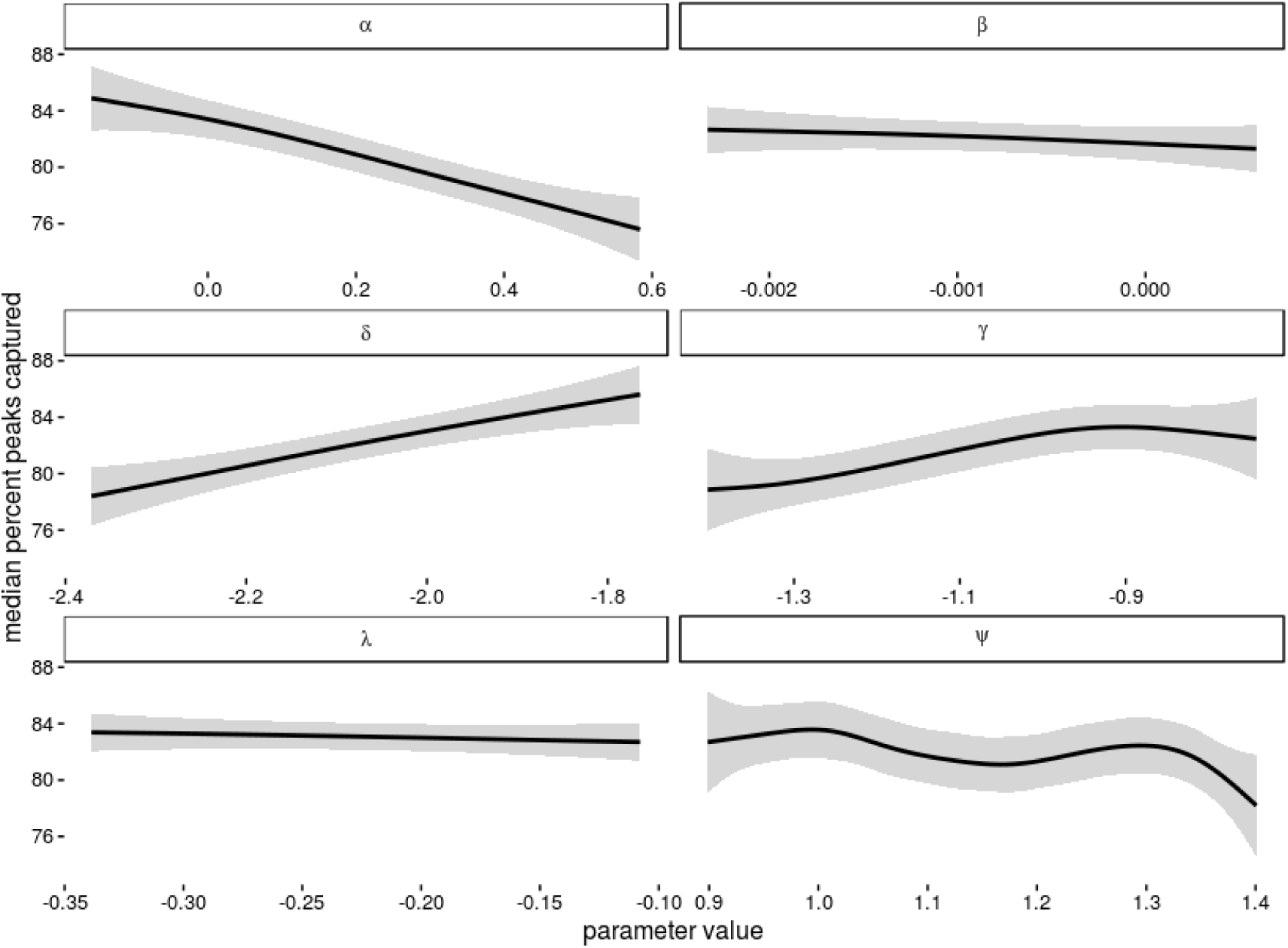
Each panel shows a different parameter used in simulation with its value on the x-axis. The smoothed conditional mean percentage of seasonal peaks included in the ALERT periods in the test data is shown by the black line. Shaded zones show the 95% confidence interval for the smoothed line.

ALERT was most sensitive to the parameters that contributed to varying levels of off-season noise and seasonality, *α*, *δ*, and *γ*. In conditions of a noisy baseline, the threshold value may activate the ALERT window too soon in the season, resulting in high *W C*_*s*_ and long *D*_*s*_. In datasets with low seasonality, the trigger may be activated too early by smaller, mini-epidemics preceding the primary seasonal outbreak. This resulted in low *P C*_*s*_ if the incidence falls below the threshold before the main peak is captured.

## DISCUSSION

The ALERT algorithm is a decision-making tool for predicting the appropriate timing of epidemic infection control measures. In this work we explained the ALERT algorithm, demonstrated how ALERT might be used on a real hospital-collected upper respiratory dataset, and used simulated datasets to test the performance measures of a training versus testing dataset.

ALERT performs best on datasets with a strong seasonal component and moderate to low off-season endemic noise. This observation is in line with the motivation behind the development of ALERT: to detect seasonal outbreaks of influenza in community healthcare settings. Across the simulations derived from the CHCO combined influenza and RSV dataset estimated parameters, ALERT was robust to modest positive and negative linear trends *β*, however, steep trends in baseline incidence will likely require a simple baseline correction before analysis. ALERT is currently designed to produce an error message if this is necessary. Similarly, ALERT was relatively unaffected by changes in *λ*, the autoregressive component, and *ψ*, the overdispersion parameter.

In the comparison of ALERT performance versus real historical periods of increased infectious disease protection implemented in a hospital setting, we found that ALERT’s performance during seasons showing multiple peaks are especially sensitive to the user’s selection of trigger value for the ALERT period. Multiple peaks during a season is common in data of mixed upper respiratory disease incidences, or even for data containing multiple strains of the same disease, where peak timing can vary. The respective heights of the peaks do not affect which one triggers the ALERT period. If a season contains two or more separate peaks, the first peak will trigger the ALERT period if the peak incidence exceeds the threshold value. If the threshold selected by the user is too high, the second peak may be missed if the nadir incidence drops below the trigger before the onset of the second peak. As ALERT was intended to apply to seasonal infectious diseases, users should proceed with caution if their data contains regular patterns of multiple peaks that they wish to capture. Likewise, users should consider recalculating potential thresholds yearly as their training dataset grows, which should help to refine the best target threshold value for their specific application.

For the training data, *D*_*s*_ and median *X*_*s*_ of the observed increased respiratory protection periods was most comparable to a *τ*_*p*_ of 10 as defined by ALERT. If *τ*_*p*_ = 10 had been used to determine increased respiratory protection periods in the testing data rather than the observed periods, ALERT would have captured a median of 5.7% more upper respiratory incidents without increasing the period duration. Both the observational method and the ALERT algorithm at *τ*_*p*_ of 10 captured more cases and fewer low weeks in the testing portion of the dataset.

The CUSUM-derived intervention periods were comparable to ALERT and the observed periods in terms of percent peaks covered, but didn’t perform as well for any of the other metrics. For this dataset, CUSUM intervention periods tended to continue past the outbreak nadir longer than necessary, so the trigger came much later than in the other two methods. This caused CUSUM to miss many cases at the beginning of the season in order to match the median duration of the other methods. CUSUM variants could likely be further optimized for use with this dataset to improve performance, as discussed in [17, 18]: however, these specialized models are difficult to implement and unlikely to be accessible to hospital epidemiologists without advanced statistical training.

Although our findings indicate that ALERT is sensitive to data seasonality and may have a tendency to trigger earlier than necessary in datasets with a noisy baseline or multiple peaks, our simulation study also shows that it is robust to a wide variety of data characteristics. Our results show promise that ALERT could be used for a variety of seasonal outbreak situations to derive a “trigger” value for the onset of epidemic periods. While we focus on influenza and RSV-derived datasets here, the simulation study results show that ALERT functions well within a broad range of dataset types, presumably including many of those derived from other seasonal diseases. Furthermore, the approach taken by ALERT and outlined in this paper has the advantage of not requiring advanced statistical training or expensive equipment. Because of these advantages, ALERT may offer an evidence-based decision-making strategy for combating infection spread in low resource settings in addition to larger hospitals that may employ an in-house epidemiologist.

Two important challenges for use of ALERT in clinical settings are 1) availability of local historical data, and 2) determining an appropriate *τ*_*s*_. While many large healthcare facilities track incidences of lab-confirmed infectious diseases, smaller facilities may not currently have the capacity for multi-year incidence recordkeeping. While there is some evidence from performance on the influenza A, influenza B and RSV combined dataset that ALERT may be used on symptom-based rather than lab-confirmed illness, this has not been studied directly to date. If only a few years of historical data is available for a site, the reliability of ALERT’s triggers may be in doubt and the threshold trigger should be re-calibrated by repeated ALERT runs as additional data becomes available. Likewise, in real data, as opposed to simulated data, we should expect that seasonality parameters may vary over time. Some of the interseason variability we observed may be due to temporal differences in these parameters. ALERT was intended for use with strongly seasonal datasets, but can accommodate some variability in these parameters. Our observations in this paper show that inconsistency resulting in occasional multiple peaks may be problematic. Likewise, differences in simulation parameters between the training and testing seasons in the simulated data would likely result in reduced ALERT performance, which is why we recommend re-calibrating the chosen threshold to new data when it becomes available. The duration of recorded historical data needed to derive useful trigger thresholds will likely vary based on seasonality and randomness characteristics of the historical data. ALERT relies on the assumption that past respiratory illness seasons will be similar to the future, which is why it performs well on seasonal data. We can observe in the more random and less seasonal simulations and in 2009 of the real data that violations of this assumption will impact ALERT’s performance. ALERT will stop with an error message if the data is not a good fit for the program, either due to unclear peak patterns or increasing or decreasing baseline. Future studies should compare the performance of ALERT to the other threshold-based methodologies for outbreak detection.

## ACKNOWLEDGEMENTS

The authors would like to thank the ResPECT study team and an anonymous reviewer for comments on earlier versions of this manuscript.

